# Visualizing Reaction Pathways via Reciprocal Space Kinetic Decomposition

**DOI:** 10.64898/2026.08.17.745189

**Authors:** Lukas Grunewald, Petra Meszaros, Sebastian Westenhoff

**Affiliations:** Department of Chemistry for Life Sciences - BMC, Biochemistry, Uppsala University; 75123 Uppsala, Sweden; Department of Cell and Molecular Biology - Molecular Biophysics, Uppsala University; 75123 Uppsala, Sweden

## Abstract

Time-resolved serial crystallography (TR-SX) has emerged as a powerful method for capturing ultrafast structural dynamics in proteins. TR-SX continues to produce remarkable studies, revealing previously unobserved transient states and providing deeper insights into processes such as drug targeting, DNA repair, and photosynthesis. However, extracting weak structural signals from noisy time-resolved datasets remains a major challenge. Robust computational methods are therefore required to isolate the signals associated with the underlying transient states. Importantly, this should be performed in reciprocal space to preserve compatibility with established downstream structure refinement workflows. Here, we introduce a framework for kinetic decomposition directly in reciprocal space that enables separation of kinetically distinct structural states. The method decomposes crystallographic data according to a predefined kinetic model, improving the recovery of weak transient signals and enhancing mechanistic interpretation from limited time-resolved datasets. We validate the framework using simulated data based on a previously published time-resolved crystallography study and demonstrate its application to a new TR-SX dataset comprising 17 time points. We show that the method separates the reciprocal space signatures of four intermediates by incorporating kinetic information from a predefined reaction model. This establishes a workflow for extracting kinetic states directly from time-resolved X-ray diffraction data that can be seamlessly integrated into existing crystallographic structure-determination pipelines.

## Introduction

Time-resolved serial crystallography (TR-SX) enables direct visualization of protein structural changes, providing atomic-level insight into dynamic biological processes across a wide range of timescales (Coquelle *et al*., 2018; Nogly *et al*., 2018; Claesson *et al*., 2020; Skopintsev *et al*., 2020; Dods *et al*., 2021, 2021; Gruhl *et al*., 2023; Safari *et al*., 2023; Glover *et al*., 2024; Shankar *et al*., 2025). In TR-SX, reactions are triggered within micrometre sized protein crystals, and the resulting changes are captured as X-ray diffraction patterns. Depending on the timescale of interest, femtosecond X-ray pulses from X-ray free-electron lasers (XFELs) can be used to investigate changes down to femtosecond timescales, while nanosecond pulses at synchrotron sources are suitable for capturing dynamics from the nanosecond to millisecond range (Chapman *et al*., 2011; Brändén and Neutze, 2021). The resulting diffraction patterns recorded at different time delays are processed to extract difference structure factors *ΔF*. Following inverse Fourier transformation, these yield difference electron density (DED) maps in real space, which provide a direct visualization of the structural changes occurring during the reaction. (Šrajer *et al*., 1996) Over the past decades, the method has been successfully applied to reveal dynamic processes such as photoactivation (Tenboer *et al*., 2014), enzyme catalysis (Butryn *et al*., 2021), and ligand binding (Barends *et al*., 2015).

Time-resolved crystallography monitors the temporal evolution of a reaction by recording diffraction data at different pump-probe delays. Except for the earliest femtosecond delay times, the measured signal is generally determined by the populations of distinct intermediate states and their structural differences relative to the reference state. Thus, the observed diffraction signal reflects the kinetics of the underlying reaction (Moffat, 2025).

Difference structure factors and the corresponding DED maps are obtained by subtracting the structure factor amplitudes of the reference state (for example the dark state) from the time-dependent diffraction data. These maps contain regions of positive and negative density arising from atomic displacements and occupancy changes (Pandey *et al*., 2020; Brändén and Neutze, 2021; De Zitter *et al*., 2022; Schmidt, 2023). Although DED maps often exhibit low signal-to-noise ratios, they provide an intuitive way to visualize structural changes. However, because the recorded diffraction pattern is generally a population-weighted average of multiple intermediate states, the resulting difference structure factors and DED maps also represent population-weighted averages of these states rather than a single structural species. Consequently, a central goal of the analysis of TR-SX data is to deconvolute difference structure factors and DED maps into the contributions of the individual intermediate states and determine their corresponding kinetics, thereby providing a mechanistic description of the reaction (Moffat, 1989, 2001).

To achieve this decomposition, we use kinetic decomposition, which has been applied widely in time-resolved fluorescence, absorption spectroscopy, and X-ray solution scattering to identify transient states and determine kinetic parameters (O’Connor, Ware and Andre, 2002; Kim *et al*., 2015; Konold *et al*., 2024). Here, the experimental data matrix *D*(*t*) can be represented as a linear combination of state-specific signals,

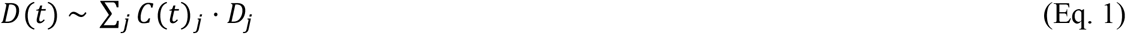

where *C_j_* describes the time-dependent population of the *j*-th state and *D_j_* its corresponding structure factor. Two or more signals can be disentangled if the kinetic profiles of the states are sufficiently different. The deconvolution also leads to averaging of the data, which is a welcome side-effect, as time-resolved structure factors usually contain small signals. To solve Eq. 1, either one of the parameters on the right must be known. If this is not the case, *C*(*t*)_*j*_ and *D_j_* can be estimated using a least square fit. The concentration profiles depend on the reaction scheme and the rate constants that connect the species. To determine these, the rate constants are iteratively optimized while recalculating *D_j_* by matrix inversion at each refinement step, thereby minimizing the residuals between the observed and re-estimated data (van Stokkum, Larsen and van Grondelle, 2004). Because the underlying reaction mechanism is generally unknown *a priori*, multiple kinetic models must be evaluated and compared. The preferred model is then selected based on its ability to describe the data accurately while maintaining minimal complexity and avoiding overfitting. However, for many studied samples information on the expected reaction scheme and its kinetic constants is already available.

As a complementary approach, singular value decomposition (SVD) is often used to analyze the information content of datasets with two independent parameters. SVD factorizes the data matrix *D* into orthogonal components according to

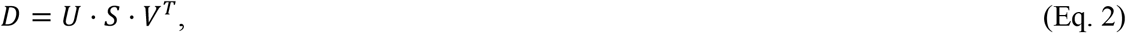

where *U* contains the temporal profiles, *V* the spectral (or spatial) patterns, and *S* the singular values ranking the significance of the components (Stewart, 1993). For time-dependent datasets, SVD is less biased than the global decomposition described above because it decomposes the data without assuming a kinetic model. Consequently, SVD is often used as a first step in global analysis. The temporal profiles (U) are subsequently fitted with an increasing number of exponential functions. Under the assumption that transitions between chemical states follow exponential kinetics, this provides an estimate of the minimum number of kinetically independent intermediate states required to describe the data.

The use of SVD for analysing time-resolved crystallographic data was first successfully demonstrated by the Moffat group (Schmidt *et al*., 2003, 2004; Rajagopal *et al*., 2004). They show that SVD (Eq. 2) can separate significant structural signals from noise in DED maps, enabling reconstruction of denoised maps and fitting of the temporal right singular vectors to kinetic models. The temporal vectors are used to identify the underlying kinetic model, while the resulting spatial components can be used to reconstruct DED maps from the intermediate states. These maps can then be converted into difference structure factors, which, when combined with the reference-state structure factors, generate extrapolated maps for crystallographic refinement of the intermediates (Schmidt *et al*., 2004). Their work established SVD as a powerful tool for the analysis of time-resolved crystallography data and demonstrated its potential for separating kinetic components in TR-SX experiments.

Building on these studies, Schotte et al. developed a global data analysis technique for time-resolved crystallography, where they applied kinetic decomposition (Eq. 1) to real space DED maps. Using a crystallographic dataset consisting of 42 time points, they refined a kinetic model against real space DED maps and recovered 4 intermediate states (Schotte *et al*., 2012). In recent TR-SX studies, decomposition methods have generally served as a supporting rather than a central role of the structural analysis. For example, Maestre-Reyna et al. assigned intermediates based on refined extrapolated structures and subsequently used SVD only to verify the proposed reaction mechanism (Maestre-Reyna *et al*., 2023). Similarly, in another study by them, they employed SVD to confirm that the experimental data were consistent with two dominant intermediates (Maestre-Reyna *et al*., 2025). In contrast, Christou et al. used both SVD and non-negative matrix factorization (NMF) on extrapolated electron density maps. The resulting components were subsequently used to guide refinement of intermediate models (Christou *et al*., 2023). Beyond classical decomposition-based approaches, machine learning methods have recently been introduced for extracting kinetic information from TR-SX data. KINNTREX, for example, utilizes a neural network to analyse time-dependent DED maps and to recover intermediate states and their associated kinetic profiles without prior assumptions about the reaction mechanism (Biener et al., 2024).

These studies demonstrate the usefulness of SVD and global analysis in TR-SX. However, so far it has only been implemented in real space. This works effectively and has the advantage that regions of interest can be selected for the kinetic devolution or SVD analysis, thereby reducing noise. However, it comes with the big disadvantage that standard crystallographic refinement routines, which typically appear downstream of the decomposition, cannot be easily used. To rectify this, the state-dependent structure factors should be obtained in reciprocal space and we propose a method to achieve this. Our method builds on obtaining the temporal profile *C*(*t*)_*j*_ (Eq. 1) by either real space refinement of a kinetic model, or from other methods, and then computing the time-dependent structure factors using this restrained temporal profile. This approach has the additional advantage that uncertainties of the reflections can be propagated, which is important for using downstream crystallographic methods for structure refinement.

## Theory

TR-SX yields difference structure factors (*ΔF_t_*) at a time delay t, with reflection indices ℎ, *k*, *l*. Each difference structure factor combines both a difference amplitude (*ΔA_t_*) and a reference phase (*φ_ref_*)

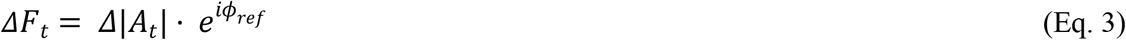

where *Δ*|*A_t_*| is defined as

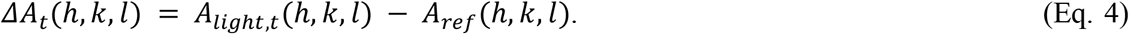

As seen in Fig. 1, the structure factor amplitudes *A_light_*, and *A_ref_* are obtained from the observed intensities (*I_obs_*). When calculating *ΔF_t_* it is important to apply thoughtful scaling, weighting, and enforcing sign consistency (De Zitter *et al*., 2022; Schmidt, 2023).

**Figure 1:**
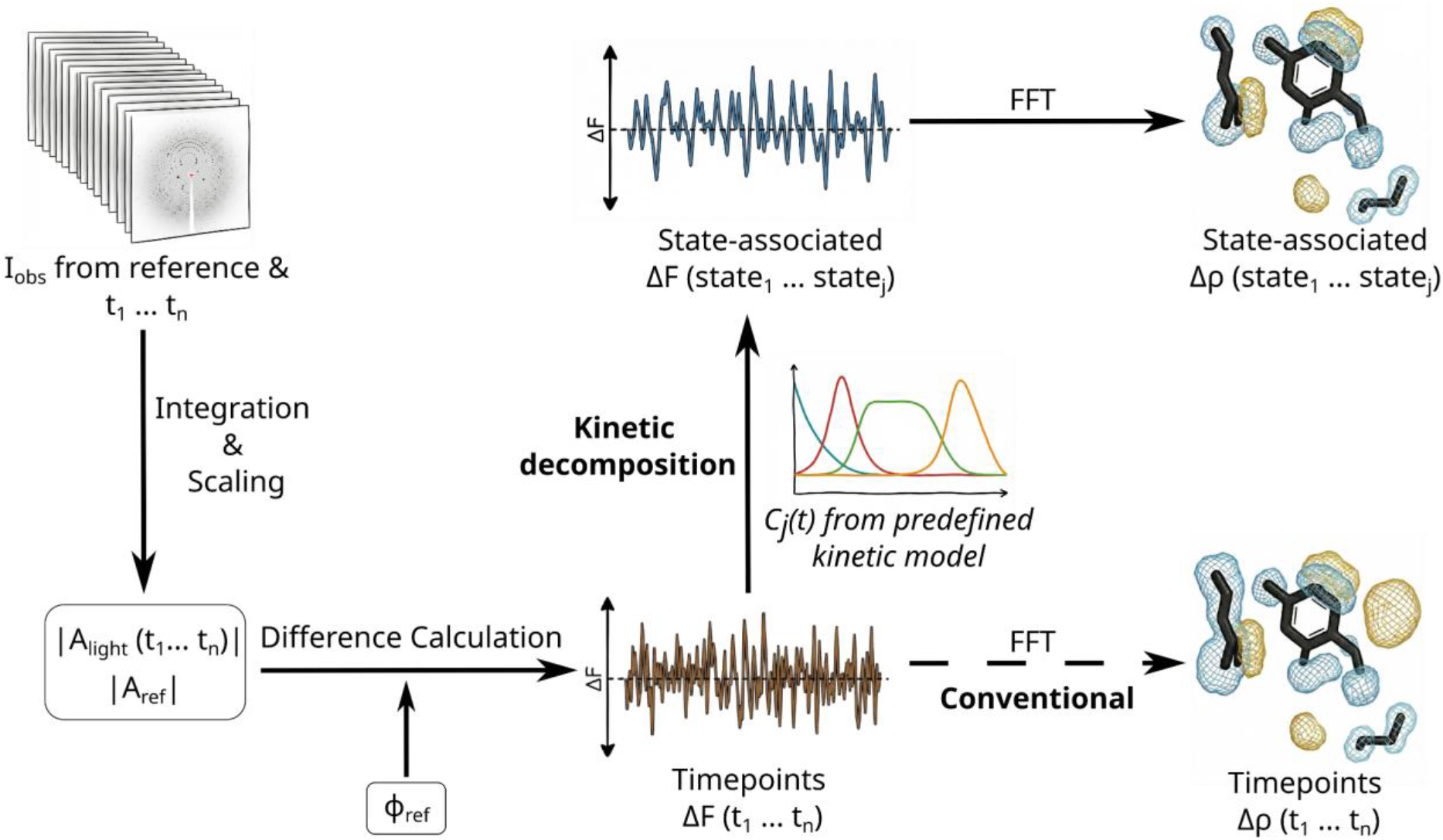
Schematic of the data analysis pipeline for time-resolved crystallography. Observed intensities (*I_obs_*) from the reference and each time point (*t*_1_ . . . *t_n_*) are integrated and scaled to obtain structure factor amplitudes (*Alig*ℎ*t*, *Aref*). Experimental difference structure factors (*ΔF*) are then calculated according to Eq. 3 using phases from the reference/dark model (*φ_ref_*). The data is processed via two streams: A conventional calculation of difference electron density maps (*Δρ*) for specific time delays via Fast Fourier Transformation (FFT) of *ΔF*, and a kinetic decomposition of the difference amplitudes *ΔA*. In the latter, time-dependent data is deconvoluted using population kinetics (*C_j_*(*t*)) from a predefined kinetic model (via SVD, spectroscopy or real space refinement), to extract state-associated structure factors. This allows for the reconstruction of electron density maps corresponding to pure structural intermediates rather than timepoints that contain mixed state amplitudes.

The central idea of a global analysis (Fig. 1) is a linear decomposition of the time-dependent dataset into states, with time-dependent concentrations. In this paper we consider reciprocal space X-ray diffraction amplitudes and assume the reference phases to be constant over the time-course of the reaction. Eq. 1 can then be written as

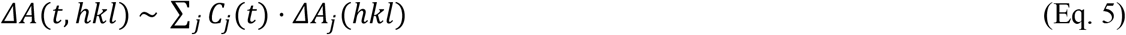

where *j* = 1 … *N* labels the *N* intermediates included in the kinetic model, *C_j_*(*t*) are time-dependent kinetic profiles and *ΔA_j_* (ℎ*kl*) are state-dependent amplitudes. Assuming exponential kinetics, the time-dependent concentrations of the states *C_j_*(*t*) can generally be described by a kinetic model formulated as a system of coupled ordinary differential equations (ODEs). In matrix notation, the time evolution of the concentration of N intermediate states *C*(*t*) = [*C*_1_(*t*), …, *C_N_*(*t*)]*^T^*

is described by

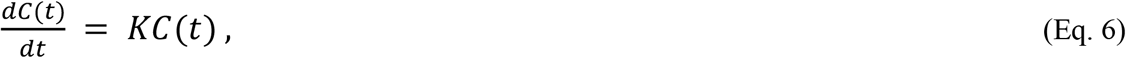

where *K* is the *N* × *N* kinetic rate matrix, constructed such that the element *K_ij_* represents the rate constant for the transition from state *j* to state *i*, and the diagonal elements *K_ii_* represents the total decay rate of state *i*. In practice, only transitions corresponding to physically plausible reaction pathways are included in the model, and all other rate constants are fixed to zero. The kinetic parameters and therefore the kinetic model of a given reaction can be determined by iteratively optimizing *K* for a given number of states to achieve the best agreement between calculated and experimental data. In practice this is often limited by the signal-to-noise level of the dataset. However, time-resolved spectroscopy, and in part, time-resolved X-ray solution scattering and diffraction datasets have been used to reliably estimate *K*. (O’Connor, Ware and Andre, 2002; van Stokkum, Larsen and van Grondelle, 2004)

For simple reaction schemes, analytical solutions can be derived directly for Eq. 6. For example, an irreversible two-state system (*A* → *B*) follows first-order kinetics and obeys classical integrated rate laws. This yields the population functions

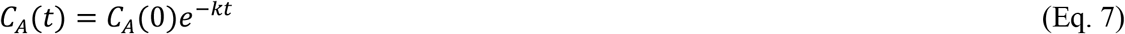

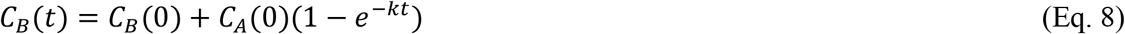

where *k* is the rate constant for conversion from *A* to *B*. Similar analytical solutions can also be obtained for certain three-state and other simple kinetic models. However, for more complex reaction networks involving multiple intermediates, branching pathways, or reversible transitions, the population profiles have to be obtained by numerically solving Eq. 6 at each iteration of the optimization procedure. (Arrhenius, 1967; Houston, 2001)

Schotte et al. applied kinetic decomposition (Eq. 5) to DED maps in real space rather than directly to reciprocal space amplitudes (Schotte *et al*., 2012). In this representation, the model becomes a decomposition of voxel intensities and takes the form of

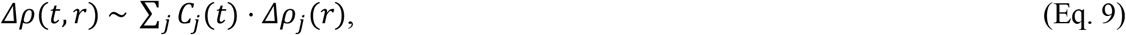

where *Δρ*(*t*, *r*) denotes the time-dependent DED maps and *Δρ_j_* (*r*) are state-specific spatial components. An advantage of the real space formulation is the ability to restrict the analysis to a defined region of interest (ROI) so that only the regions with actual signals are included in the optimization procedure, which often is important for optimization of the kinetic coefficients *C_j_*(*t*). In reciprocal space, such spatial localization is not possible, as the data are distributed over reflections rather than physical coordinates. This makes the estimation of *C_j_*(*t*) more sensitive to noise and in practice it is practically impossible (Schotte *et al*., 2012). Thus, we use predetermined kinetic models, for example by analysing complementary spectroscopy data or using real space data and ROIs to estimate the concentration profiles in Eq. 5.

One advantage of performing the decomposition in reciprocal space is that the experimental uncertainties can be retained and consistently propagated into the state dependent amplitudes, which would be lost if done in real space. This is done by propagating the uncertainties *σ* of the measured amplitudes into the kinetic basis amplitudes, such that the variance of each basis amplitude is given by a weighted sum over time

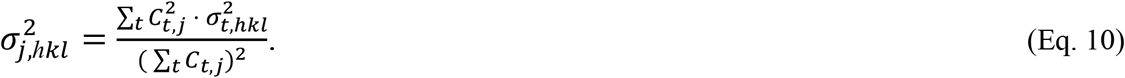

The propagation of experimental uncertainties from the time-dependent observations into the kinetic states is important, because they enable the statistical down-weighting of reflections where the state dependent structure factor is dominated by noise using the per-state reflection weights *ω* by the procedure of Ursby and Bourgeois (Ursby and Bourgeois, 1997)

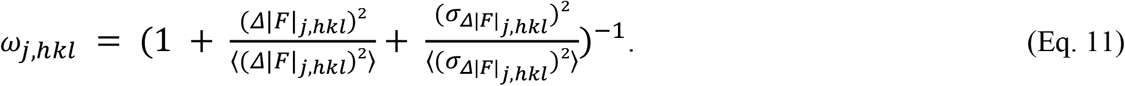

The weighting scheme depends on the magnitude of the individual difference structure factor, *Δ*|*F*|_*j,hkl*_, and its associated uncertainty, *σ_Δ|F|j,hkl_*, relative to their average values across the dataset. The difference amplitudes can then be normalized by the average reflection weight

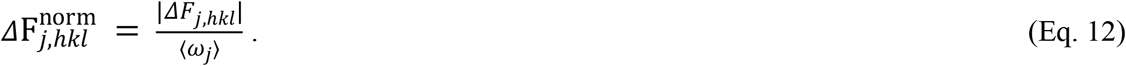

Afterwards, these normalized difference amplitudes can be combined with the reflection- specific weights *ω_j,hkl_* during the generation of DED maps, as well as for the calculation of extrapolated structure factors which are required for structural refinement. The extrapolated structure factors can then be calculated by

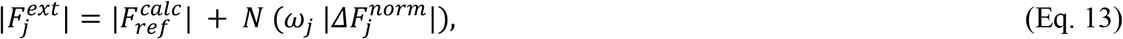

where N is an extrapolation factor which corresponds to the photoactivation yield 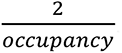.

This weighting approach ensures that the subsequent structural refinement is not driven by noise amplified during the extrapolation process.

## Results

### 1. Validation on simulated data

First, we benchmark the kinetic deconvolution algorithm (Eq. 5) on simulated difference structure factor amplitudes. To this end, we constructed 17 time points (see Table 1) containing different concentrations of four known structural intermediates of the photoactive yellow protein (PYP) which were resolved by Schotte et al. (Schotte et al., 2012). These timepoints were generated by calculating structure factor amplitudes from the structure of each intermediate (*pR*_0_, *pR*_1_, *pR*_2_ and *pB*_0_ with the PDB ID codes 4B9O, 4BBT, 4BBU, and 4BBV respectively), adding varying degrees of resolution-dependent noise, and mixing them according to predefined concentrations at each delay (Fig. 2A, see methods). To facilitate visualization, the structure factors were inverse Fourier transformed into real space (Fig. 2B,D,F).

**Figure 2:**
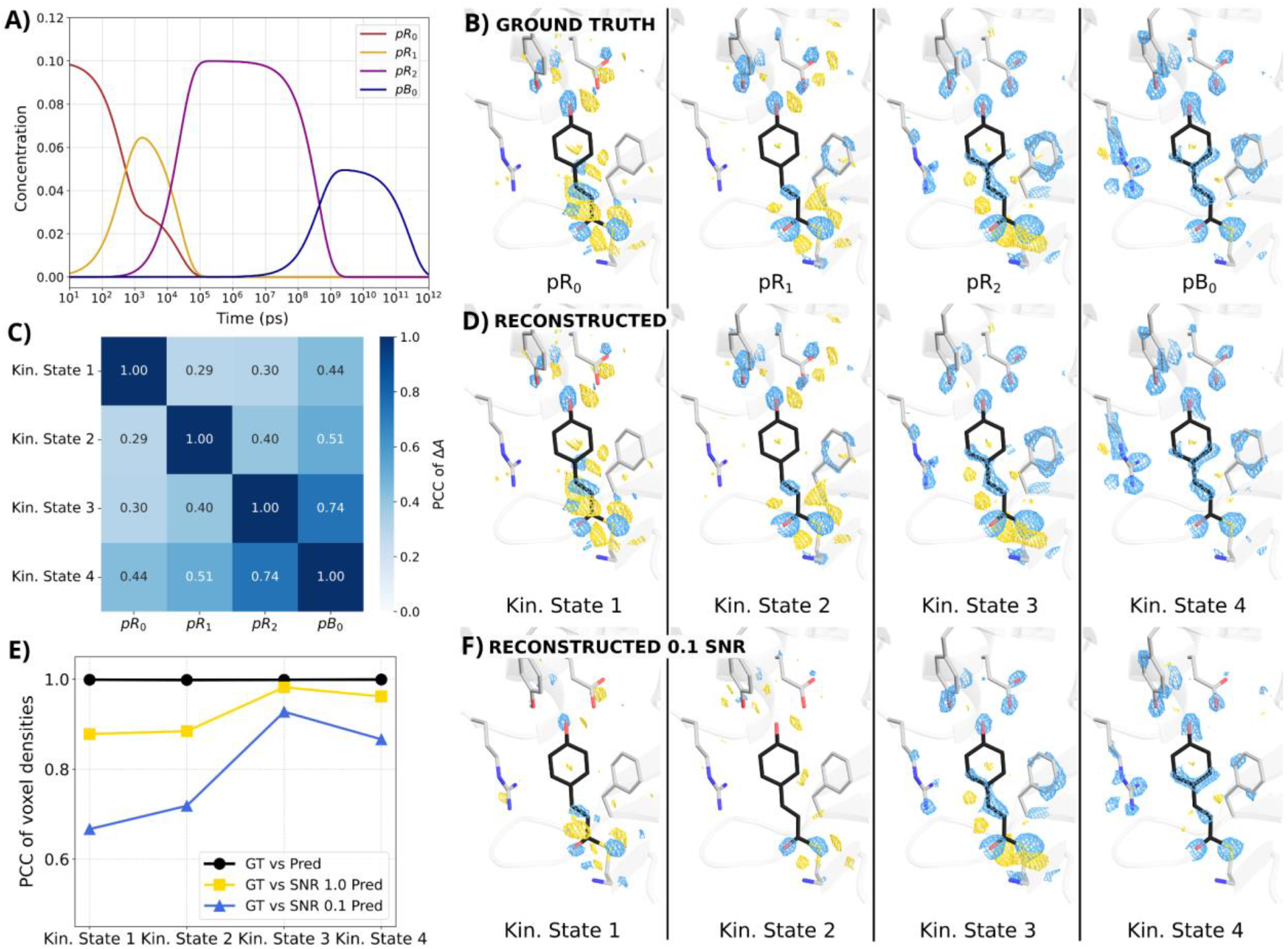
Validation of Kinetic Mode Reconstruction against ground truth “PYP” data. A) The population dynamics of the four modelled states over time, indicating the assumed 10% quantum yield. B) Ground truth difference electron density maps contoured at 3*σ* for the four simulated states. Gold meshes indicate positive difference density and blue meshes indicate negative difference density. C) Correlation matrix showing the Pearson Correlation Coefficient (PCC) between the reconstructed and ground truth difference amplitudes (*ΔA*) in Fourier space. High diagonal values indicate accurate state recovery. D) The corresponding kinetic modes at 3*σ* reconstructed using the deconvolution algorithm (Eq. 5) on the ground truth data. E) PCC calculated in real space, assessing the robustness of the reconstruction against noise. The plot shows the correlation between the ground truth (GT) maps and predictions derived from: the clean input data (Pred, black circles), input data with a Signal-to-Noise Ratio (SNR) of 1.0 (SNR 1.0 Pred, yellow squares), and input data with an SNR of 0.1 (SNR 0.1 Pred, blue triangles). F) Reconstructed SNR 0.1 difference electron density maps contoured at 3*σ*, which shows the robustness of the deconvolution algorithm against noise.

**Table 1:** Kinetic deconvolution improves the signal-to-noise ratio of difference structure factor amplitudes. * The values for individual timepoints were calculated first by determining 〈*F_hkl_*/π_*hkl*_〉 for each timepoint up to 100 ps and then averaging these values across all timepoints.

| | $\langle \Delta F_{hkl}/\sigma_{hkl} \rangle$ | % $> 1\sigma$ | % $> 2\sigma$ | % $> 3\sigma$ |
| --- | --- | --- | --- | --- |
| Kin. State 1 | 5.54 | 87.4% | 75.3% | 63.8% |
| Kin. State 2 | 4.85 | 84.9% | 70.2% | 57.0% |
| Kin. State 3 | 4.65 | 84.1% | 68.8% | 55.1% |
| Timepoints* | $\sim 3.13$ | $\sim 76\%$ | $\sim 57\%$ | $\sim 37\%$ |

After generating the simulated mixed amplitudes, we applied the kinetic deconvolution defined by Eq. 5 using the known concentration profiles (Fig. 2A) to recover the state dependent difference amplitudes. In the absence of adding extra noise, the algorithm reconstructs the ground truth states with numerical precision, yielding Pearson Correlation Coefficients (PCCs) of 1.0 for all states (Fig. 2B), global R-values of zero (see Methods), and visually indistinguishable electron-density features between the true and recovered maps (Fig. 2B,D). This verifies the correctness of the implementation.

We then continued to assess the algorithmic robustness to noise by analysing residuals in Fourier space (see methods) and PCCs in real space after adding different levels of noise to the mixed time-point amplitudes (Fig. 2E). We tested two noise regimes: an SNR of 1.0 and an extreme SNR of 0.1. Notably, the uncertainties used in the weighting scheme were not adjusted to match this added noise, creating a challenging scenario where the algorithm must handle errors underestimated by the input model.

The PCC analysis in real space demonstrates strong robustness of the kinetic deconvolution, maintaining PCC values above 0.85 at SNR 1.0 for all recovered states. States 3 and 4 exhibit particularly high robustness to noise, which can be attributed to their temporal isolation in the concentration profiles (Fig. 2A), where they dominate the population over distinct time windows. Additionally, the DED signals of states 3 and 4 are stronger due to larger structural changes of the nearby amino acid side chains and the chromophore. In contrast, states 1 and 2 exhibit greater sensitivity to noise, likely due to their substantial temporal overlap, which reduces the separability of the kinetic decomposition, combined with weaker DED signals. Nevertheless, at the highest noise level tested (SNR 0.1), the reconstructed real space maps remain interpretable, although weaker density features are obscured by noise (Fig. 2F). Importantly, the characteristic structural changes surrounding the chromophore are retained.

This benchmark confirms that the algorithm performs reliably, i.e. that it recovers the difference structure factors for states that partially overlap in time. Performance remains robust at the realistic noise level of SNR 1.0, supporting the applicability of the method to real experimental datasets.

### 2. Application to experimental data

Having established the performance of the approach on simulated data, we applied the kinetic deconvolution to experimental time-resolved crystallographic data for a bacterial phytochrome protein, where a light-induced reaction was triggered and monitored from femto- to microsecond times scales over 17 timepoints. A full analysis of the structural conclusions is described in a separate paper (Mészáros et al., 2026). Here, we use the dataset to investigate the kinetic decomposition only.

Applying the deconvolution in reciprocal space requires that the data matrix has to be defined over a consistent set of reflections across all timepoints. This requirement is not a fundamental limitation of the kinetic deconvolution itself, but a consequence of the current implementation: in real space, every difference map is defined on the same voxel grid in a ROI, while in reciprocal space, each timepoint’s reflections can contain a different subset of measured Miller indices. The *hkl* indices must therefore be matched across timepoints before we apply the deconvolution. In addition, difference structure factor amplitudes must be adjusted to the dark-state phases, with amplitudes sign-flipped for reflections exhibiting 180° phase differences.

Mathematically, the analysis only requires a number of timepoints sharing a common set of reflections that is equal to or greater than the number of underlying kinetic states. In practice, however, we found that operating close to this minimum resulted in increased numerical instability. We therefore imposed a stricter, empirically determined reflection coverage threshold. Depending on the applied threshold, approximately 10,000 to 50,000 reflections (as seen in Fig. 3) were excluded from the global analysis. Restricting the analysis to reflections present at all 17 timepoints removes a substantial fraction of the data (∼40%). In contrast, retaining reflections observed in at least 12 of the 17 timepoints (a reflection coverage threshold of ∼70%) produces maps with similarly high quality and correlation coefficients (Fig. 3C), while preserving substantially more reflections.

**Figure 3:**
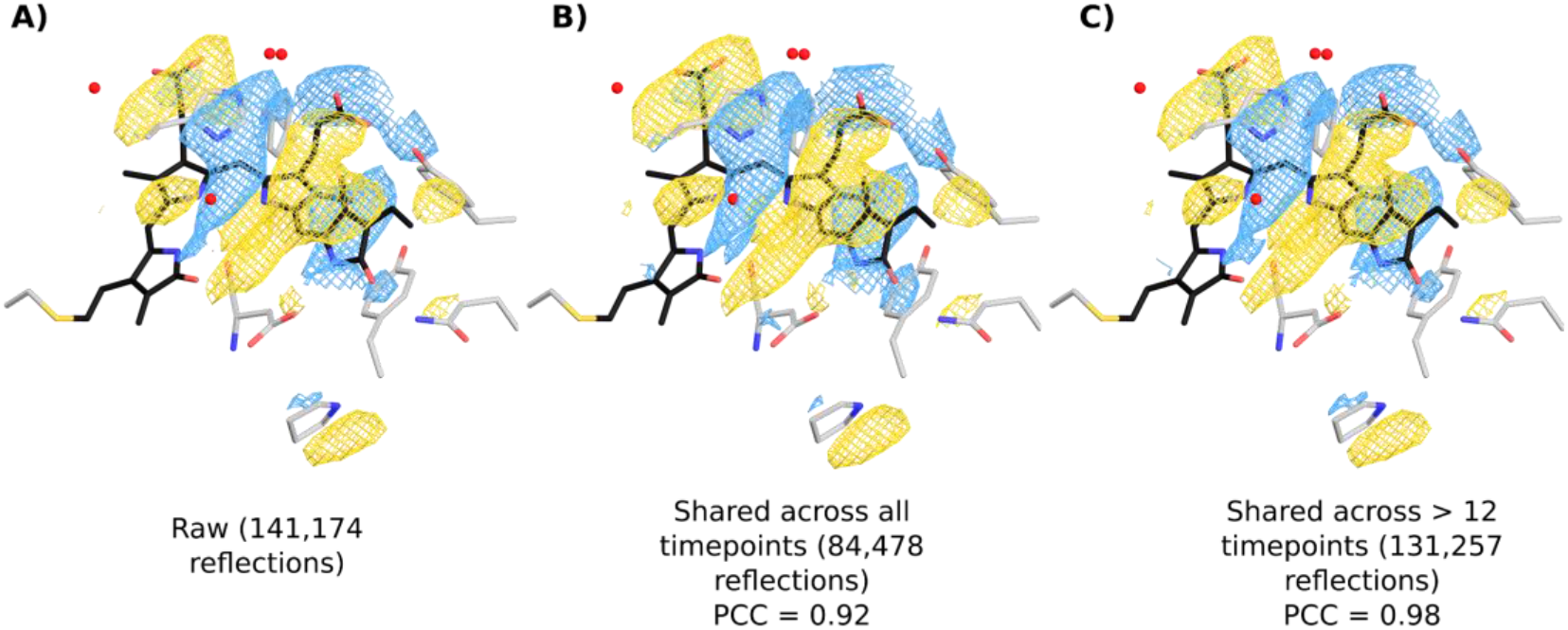
Effect of reflection filtering on DED maps. A) Raw difference electron density map including all measured reflections (141,174). B) Map after retaining only reflections present at all delay times (84,478), showing slightly reduced map detail but high Pearson correlation (PCC = 0.92). C) Map using a relaxed threshold of reflections observed in more than 12 out of 17 timepoints (131,257), maintaining high map quality and improved correlation (PCC = 0.98). Blue and yellow indicate positive and negative difference densities, respectively.

Following these preprocessing steps, we applied the kinetic deconvolution to the experimental data (Fig. 4, 4A). As discussed in the introduction, we restricted the kinetic deconvolution analysis to timepoints from 200 fs onward, excluding the earliest, impulsive regime in which the observed signals likely reflect coherent atomic motion rather than a mixture of intermediate states. A four-state kinetic model derived from prior spectroscopic measurements and refinement of the parameters using real space DED maps (Eq. 9, Fig. 4B) was used to describe the temporal evolution of the photoreaction. Applying this model to the structure factor amplitudes of 17 timepoints yielded reconstructed DED maps that reproduce the structural features observed in the raw experimental data (Fig. 4C). The kinetic deconvolution effectively separates the time-dependent series into four states.

**Figure 4:**
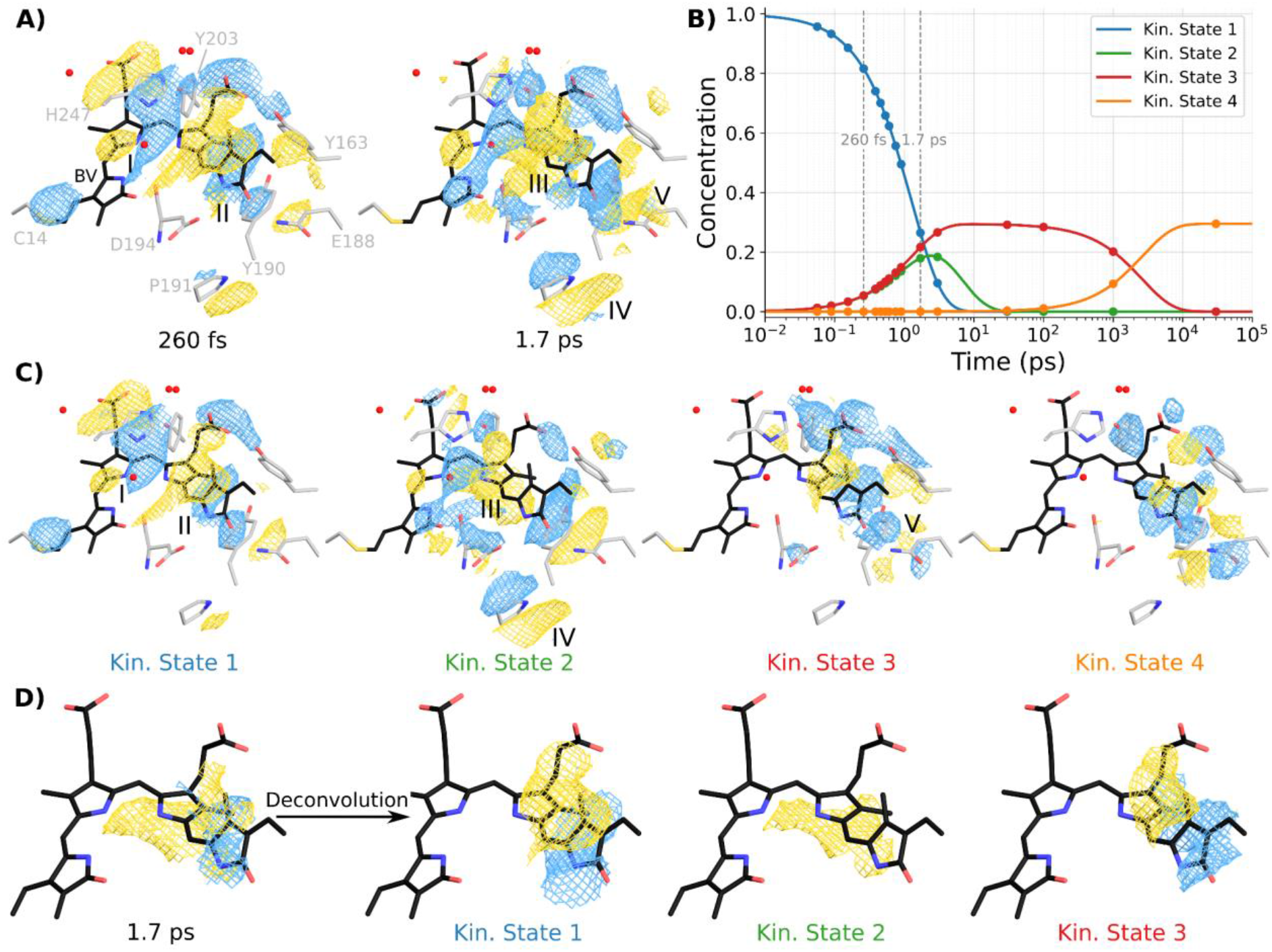
Experimental deconvolution and global kinetic analysis of phytochrome TR-SFX data. A) Example experimental difference electron density maps observed at 260 fs and 1.7 ps delay times for the phytochrome. Gold and blue isosurfaces indicate positive and negative difference densities, respectively. The DED maps shown are averaged over all eight monomers in the asymmetric unit. B) Population kinetics of the four recovered states derived from global analysis. The plot displays the concentration of each kinetic mode (1–4) over time, with vertical dashed lines marking the 260 fs and 1.7 ps time points shown in Panel A. C) Reconstructed species-associated difference maps of the four kinetic modes. These DED maps represent the deconvolved intermediate states, separating specific features. D) Focus on the chromophore-localized DED maps at the 1.7 ps timepoint.

Visual inspection of the resulting state-dependent DED maps (Fig. 4C) shows that all four states carry different signatures: a negative peak at feature I in states 1 and 2; a negative/positive pair at the biliverdin (BV) chromophore isomerization site, feature II; a positive peak beneath the BV, feature III; a negative/positive pair at feature IV specific to state 2; and a strong signal at feature V shared by states 3 and 4. The feature IV signal is particularly clear of the deconvolution’s power, as it is essentially absent from states 1 and 3 and appears only in state 2.

While the earliest and latest time points are dominated by a single kinetic state, intermediate delays, such as 1.7 ps, contain substantial contributions from multiple states (Fig. 4B). We therefore focus on chromophore-specific changes at the 1.7 ps time point (Fig. 4D). Inspection of kinetic states 1–3 reveals that difference density around the D-ring is confined to states 1 and 3, whereas state 2 exhibits a pronounced positive density feature beneath the chromophore. Thus, the kinetic deconvolution filters out the density of state 2, which would otherwise be hidden by the dominating signals of states 1 and 3. Moreover, if the mixed signal at 1.7 ps were used directly for structure refinement, these distinct structural signatures would be averaged together, obscuring the underlying state-specific rearrangements. In contrast, kinetic deconvolution separates the signal into three clean, individually interpretable states that can each be refined independently.

Importantly, the deconvolution in Fourier space enables propagation of experimental uncertainties (see theory). This allows us to compute difference structure factors and to use standard refinement of structures against the deconvoluted data based on extrapolated structure factors. We have confirmed that structures can be refined against the state-decomposed data, which will be reported in a separate paper (Mészáros *et al*., 2026)

As a consequence of the kinetic decomposition, each state is effectively averaged over several time points. This leads to an increase in SNR for the difference structure factors. For example the mean *ΔF*/*σ* for state 1-3 is roughly 1.5× higher than for the individual timepoints from which they are derived, with more than 55% of reflections in each state exceeding *ΔF*/*σ* > 3, compared to an average of only ∼37% across the individual timepoints up to 100 ps (see Table 1).

Validation of the deconvolved state-dependent difference maps is important, but difficult to achieve because ground truth, state-dependent structure factors are not available for experimental datasets. We therefore assess the quality of the decomposition by comparison of the species-dependent *ΔF_j_* to the time-dependent, measured experimental structure factors *ΔF*. This is quantified using a PCC analysis in real space (Fig. 5) between the real space DED maps calculated for the kinetic states and each experimental timepoint. We note that this analysis does not confirm the correctness of the kinetic model, as the concentration profiles are fixed inputs rather than fitted parameters. Therefore, the PCC analysis serves to verify that the decomposition has been implemented correctly and behaves as expected and provides a diagnostic check against the underlying concentration profiles. High correlations at early delay times (200–400 fs) with state 1 indicate that this state dominates the signal in this regime, consistent with the global kinetics in Fig. 4B. Similarly, later timepoints show good correlation with corresponding kinetic modes, reflecting the expected temporal evolution of the system. Even for state 2, which exists transiently with a maximum at around 1 ps, we find corresponding kinetics in the PCC plot (Fig. 5). Overall, the PCC analysis supports that the deconvolution uses a consistent kinetic model that reproduces the experimental data and correctly separates the underlying states.

**Figure 5:**
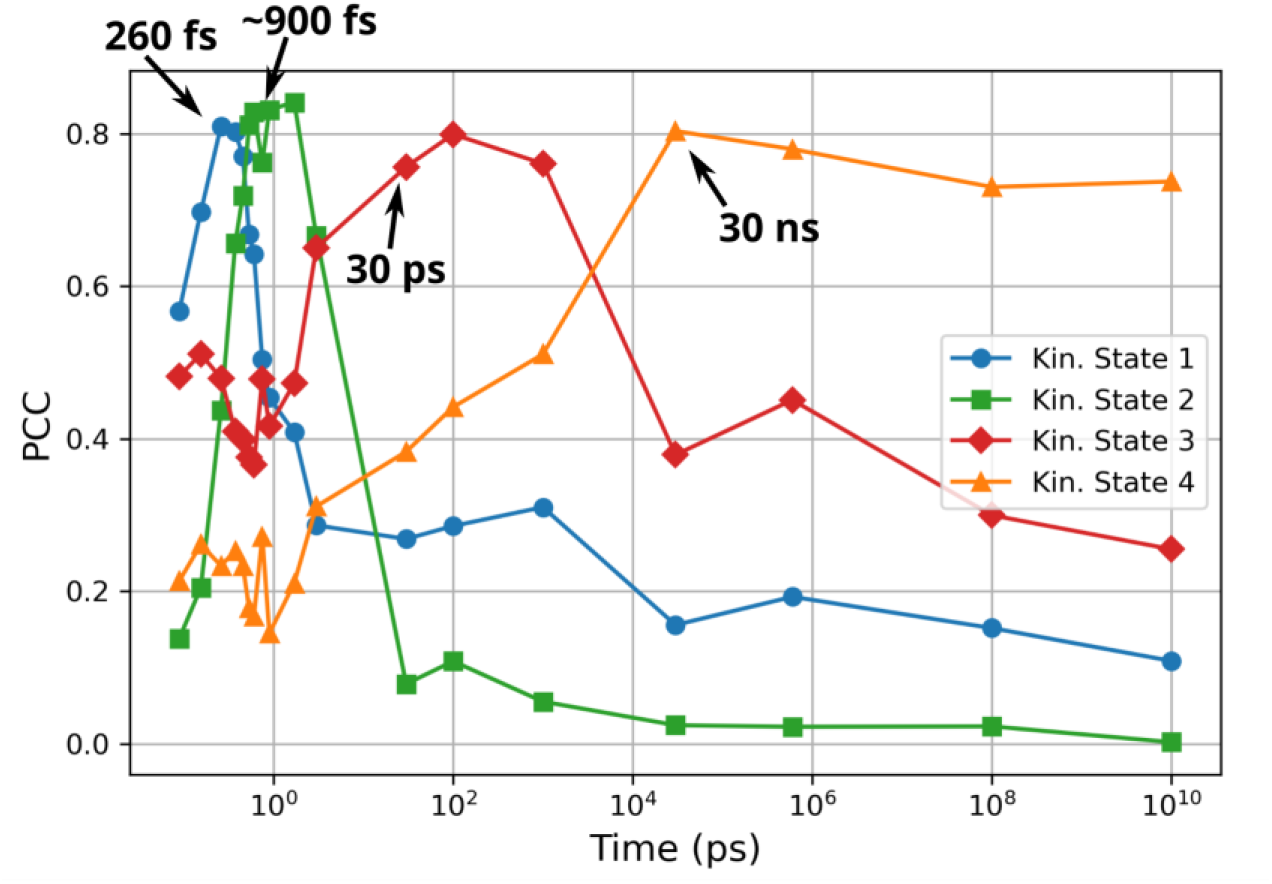
Real space Pearson correlation coefficient (PCC) analysis of kinetic modes. PCC within the binding pocket shows rapid initial transitions, with Kinetic Mode 1 peaking before 400 fs and Mode 2 peaking near 1.7 ps. This progression aligns with the kinetic rates observed in Fig. 4B.

## Discussion

Here, we report a framework for extracting structure factors of kinetic states from time-resolved crystallographic data via reciprocal space deconvolution. This approach exploits the linear independence of structure factors in Fourier space to resolve overlapping structural signals. This adds a new method to the growing toolbox for processing, analysing, and interpreting time resolved crystallographic data (White *et al*., 2012; Dalton, Greisman and Hekstra, 2022; De Zitter *et al*., 2022; Brookner and Hekstra, 2024; Fadini *et al*., 2025; Perrett *et al*., 2025). Together, these approaches establish increasingly rigorous workflows for TR-SX.

A key aspect of the reciprocal space deconvolution is that it operates directly on experimentally measured, noise-containing difference structure factors. This is different from previous published studies, where deconvolution or SVD was performed in real space. (Schmidt et a*l.*, 2003, 2004; Rajagopal *et al*., 2004; Schotte *et al*., 2012). While the data are inherently noisy, the linearity of the decomposition ensures that noise is propagated without introducing additional bias or amplification during reconstruction.

An important advantage of performing the deconvolution in reciprocal space is the preservation of the uncertainties of individual structure factors. Since the deconvolution operates directly on difference structure factors, these uncertainties can be propagated throughout the analysis. This enables the calculation of extrapolated structure factors and the application of established crystallographic procedures, including structural refinement in reciprocal space. In contrast, propagation of uncertainties through deconvolution in real space is not possible, which restricts subsequent structural refinement to real space analysis, which is inferior to the reciprocal space refinements. Reciprocal space deconvolution can therefore be integrated naturally into existing crystallographic workflows.

In the present framework, the temporal evolution of the states is assumed to be known from independent measurements and is used to decompose the experimental data into state-specific contributions. In contrast, real space kinetic refinement approaches can, in principle, optimize rate constants directly against the crystallographic data, allowing simultaneous refinement of both structural states and kinetics. We note that one can use these approaches in a complementary way: One could refine the rates in the kinetic model in real space, using a region of interest, and then deconvolute in reciprocal space, thereby propagating the uncertainties.

Finally, kinetic deconvolution relies on the assumption that the time dependent signals can be described as a linear combination of structurally distinct states with well-defined populations. This approximation is reasonable for most delay times, however, at very early time delays, structural changes are likely dominated by coherent atomic motions within one or more states rather than population transfer between discrete states. At these short timescales, the structures have not yet reached their equilibrated form, and the atomic positions evolve continuously with time. In this regime, the population based description of Eq. 5 does not fully apply. Moreover, the transition between the regimes is not clearly defined in time as it depends on the type of motion and potential transition times between states. A rigorous kinetic decomposition that simultaneously accounts for both population dynamics and continuous structural evolution would be highly desirable but requires more future work.

## Practical guidelines

We finish by providing practical guidance for application of the method, which we have made available via GitHub (https://github.com/Westenhoff-Lab/fourier-kin-dec). When kinetic decomposition is planned as part of the data analysis, it is advantageous to prioritize acquiring more timepoints with fewer diffraction patterns each, rather than fewer timepoints with higher pattern counts. Since kinetic decomposition averages out noise effectively, denser sampling in time yields more temporal information for the model. After the experiment, we suggest computing DED maps as usual for visual inspection, however, if several states are expected to overlap temporally, the data must be kinetically decomposed to obtain the structure factors of the individual intermediates. This requires prior information, e.g. spectroscopic data or doing a SVD analysis on the real space DED maps. The specifics will depend on the system under investigation. For studies covering the earliest femtosecond timescales, we suggest deconvolving data starting between 200–500 fs. The transition from the impulsive regime to the regime where the discrete-state approximation applies is not well defined and depends in a complex way on the available timepoints, time resolution, and photophysics of the system. A practical approach is to check whether the resulting state-dependent DED maps vary with the chosen cutoff. Finally, after kinetic decomposition it is important to verify that the resulting DED maps actually differ between states. If they do not, this likely indicates that a proposed state or its temporal profile was not specified correctly, and the kinetic model should be revisited.

## Conclusion

In summary, reciprocal space kinetic deconvolution provides a framework for separating overlapping structural signals in time-resolved crystallographic data while preserving the experimental information required for direct downstream crystallographic analysis. The method complements existing real space approaches and offers a practical route for the structural characterization of transient intermediates in complex reaction pathways.

## Methods

### Simulation of time-resolved data

To benchmark the deconvolution algorithm against a realistic dataset which shows overlapping states in the timepoints, we used the resolved photocycle intermediates of Photoactive Yellow Protein (PYP) previously resolved by Schotte et al. (Schotte *et al*., 2012). Their study reported a total of 42 time delays, from which they calculated a kinetic model to resolve four structural intermediates named *pR*_0_, *pR*_1_, *pR*_2_ and *pB*_0_ (PDB ID codes 4B90, 4BBT, 4BBU, and 4BBV respectively). The dark state structure was adopted from Yamaguchi et al. (Yamaguchi et al., 2009, PDB ID code 2ZOH).

Ground truth structure factor amplitudes *F_calc,j_* were simulated from each state j using phenix.fmodel with a resolution range of 57.91-1.80 (Liebschner *et al*., 2019). To mimic experimental error, resolution dependent noise was added to these amplitudes. We applied a noise model where the target Signal-to-Noise Ratio (SNR) decays linearly as a function of the squared inverse resolution (*s*² = 1/*d*²). The noise level was roughly calibrated to yield a mean SNR of 30.0 at the lowest resolution shell (57.91-3.88), decreasing to an SNR of ∼1.0 at the highest resolution shell (1.87-1.80).

For each reflection, a standard error (*σ*) was derived from the target SNR, and the final simulated amplitude was generated by adding random Gaussian noise sampled from *N*(0, *σ*):

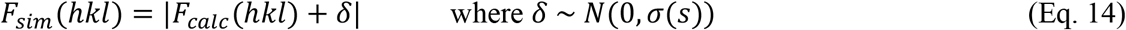

This ensures that high-resolution data is appropriately down-weighted during analysis, replicating the statistical quality typical of TR-SX experiments.

From these four states, difference structure factors *ΔF_sim_*(*hkl*) and their uncertainties were calculated in CCP4 (Agirre et al., 2023) according to Eq. 3 and 4 after scaling, weighting and forcing sign consistency (Schmidt, 2023). To avoid phase inconsistencies during mixing, all difference structure factors were expressed relative to a common dark-state phase origin prior to mixing: reflections with a phase shift of 180° relative to the dark state were reassigned the dark-state phase and the sign of the amplitude was flipped accordingly. This allows the mixed dataset to be constructed as a simple linear combination of signed scalar amplitudes rather than requiring explicit phase tracking. After deconvolution, the signs were reverted to restore the standard convention of positive amplitudes in the final maps.

To generate the dataset of timepoints at different time delays, we roughly used the populations from Schotte et al.’s population plot (see Fig. 4B in Schotte et al., 2012). The final occupancies are listed in Table 2.

**Table 2.** Time-dependent occupancies used for the simulated dataset. The populations for each structural intermediate (pR0, pR1, pR2, pB0) were defined at various time delays to mimic a realistic photocycle progression.

| Time Delay | pR0 (Red) | pR1 (Yellow) | pR2 (Purple) | pB0 (Blue) |
| --- | --- | --- | --- | --- |
| **100 ps** | 0.080 | 0.010 | 0.000 | 0.000 |
| **300 ps** | 0.060 | 0.025 | 0.000 | 0.000 |
| **500 ps** | 0.045 | 0.040 | 0.000 | 0.000 |
| **1 ns** | 0.035 | 0.050 | 0.005 | 0.000 |
| **2 ns** | 0.030 | 0.060 | 0.010 | 0.000 |
| **3 ns** | 0.028 | 0.065 | 0.015 | 0.000 |
| **5 ns** | 0.025 | 0.060 | 0.018 | 0.000 |
| **7 ns** | 0.023 | 0.050 | 0.020 | 0.000 |
| **10 ns** | 0.020 | 0.040 | 0.040 | 0.000 |
| **30 ns** | 0.010 | 0.020 | 0.090 | 0.000 |
| **50 ns** | 0.005 | 0.010 | 0.100 | 0.000 |
| **1 μs** | 0.000 | 0.000 | 0.105 | 0.000 |
| **10 μs** | 0.000 | 0.000 | 0.100 | 0.005 |
| **100 μs** | 0.000 | 0.000 | 0.080 | 0.015 |
| **1 ms** | 0.000 | 0.000 | 0.005 | 0.050 |
| **10 ms** | 0.000 | 0.000 | 0.000 | 0.040 |
| **100 ms** | 0.000 | 0.000 | 0.000 | 0.030 |

### Robustness Analysis and Noise Injection

To evaluate the stability of the kinetic deconvolution algorithm under varying noise conditions, we generated additional datasets by injecting Gaussian noise directly into the mixed time-point amplitudes *A_mixed_*. Unlike the resolution dependent measurement noise added during the state dependent simulations (see Eq. 14), which results in higher signal to noise ratios at low resolution and lower signal to noise ratios at high resolution, this step applies uniform noise across all reflections, yielding a constant signal to noise ratio independent of resolution. This represents a severe test condition in which even low resolution reflections are strongly affected by noise.

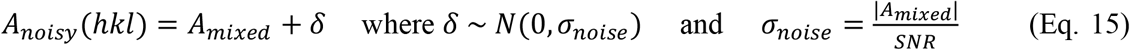

The results of this robustness analysis are shown in Fig. 2E (real space) and below in Fig. 6 (reciprocal space, residuals between the ground truth and predicted difference structure factor amplitudes), demonstrating that the algorithm recovers state information with high correlation (PCC > 0.8 for most cases in real space). As seen in Fig. 6, clean reconstructions yield perfect global R-factors of 0.0 with Correlation Coefficients of 1.0. While the introduction of noise increases the global R-factors to a range of 0.35–0.61 (SNR 1.0) and 3.7-5.05 (SNR 0.1), the correlation coefficients remain robust between 0.70–0.85 (SNR 1.0) and 0.56-0.69 (SNR 0.1). These values indicate that despite the high noise level, the deconvolution algorithm maintains the ability to recover the underlying ground truth signal in reciprocal space with moderate to high confidence.

**Figure 6:**
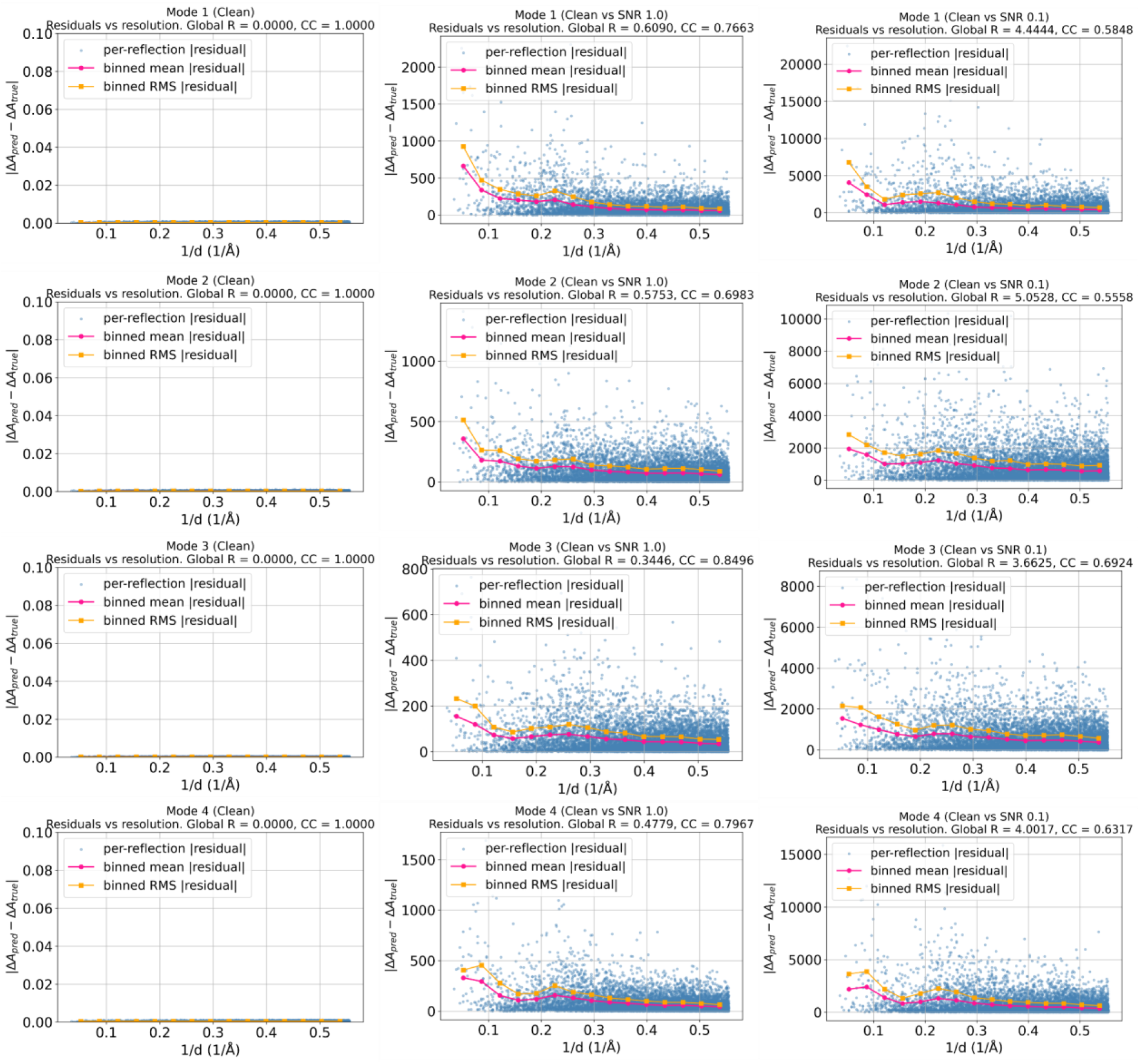
Fourier space deconvolution accuracy across varying noise levels. Rows represent reconstructed kinetic modes 1–4. Columns correspond to input data quality: clean (left), Signal-to-Noise Ratio (SNR) 1.0 (middle), and SNR 0.1 (right). Plots map difference amplitude residuals (|*ΔA_pred_* − *ΔA_true_*|) against resolution (1/*d*). Blue dots indicate per-reflection residuals; magenta lines mark binned mean residuals; orange lines denote binned RMS residuals.

### Experimental data preprocessing

Difference structure factor amplitudes were treated relative to the dark-state phase and sign-flipped where necessary. Before kinetic deconvolution, datasets were filtered to yield a consistent set of Miller indices across all timepoints. For the final deconvolution shown in Fig. 4, a threshold of at least 12 indices shared between the timepoints was chosen (see Results). After deconvolution, the signs were reverted to restore the standard convention of positive amplitudes in the final state dependent maps.

The time-dependent concentration profiles were derived from time-resolved infrared spectroscopy and optimized in real space using a four-state kinetic model:

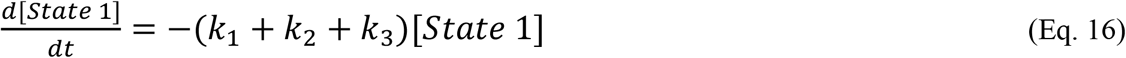

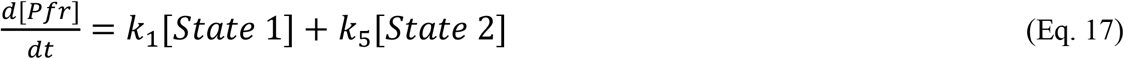

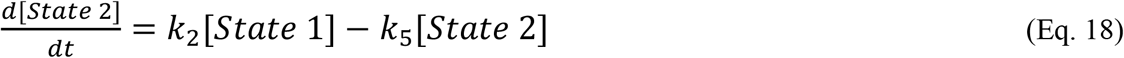

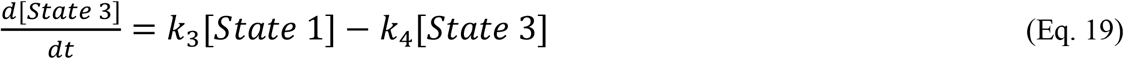

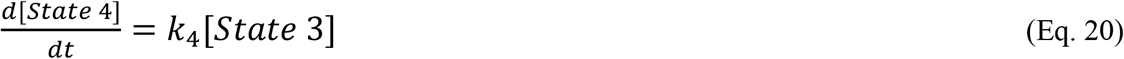

where Pfr denotes the dark-state population, included in the rate equations but not part of the concentration profile *C_j_*(*t*) used for the four recovered kinetic states. The fitted rate constants were k1 = 0.3204 ps⁻¹, k2 = 0.2304 ps⁻¹, k3 = 0.2304 ps⁻¹, k4 = 0.00038 ps⁻¹, and k5 = 0.1881 ps⁻¹.

### Kinetic deconvolution of experimental data

The basis amplitudes *M_j_*(*hkl*) were then obtained by solving the linear least squares problem for Eq. 5 by:were then obtained

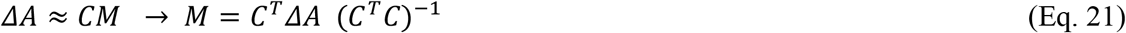

